# Ovarian hormonal state at exercise initiation interacts with nigrostriatal circuitry to determine long-term voluntary exercise behavior

**DOI:** 10.64898/2026.06.17.732957

**Authors:** Margaret K Tanner, Kamryn M Korth, Alyssa A Hohorst, Juliet R Freund, Jessica D Westerman, Carolina Sanchez Mendoza, Benjamin N Greenwood

## Abstract

Despite the well-established health benefits of exercise, adherence to physical activity remains low, highlighting the need to identify biological factors that regulate the initiation and persistence of exercise behavior. Here, we tested whether ovarian hormone state at the onset of voluntary wheel running (VWR) influences both immediate and long-term exercise behavior in female rats. Females that initiated VWR during proestrus (Pro) ran farther, spent more time running, and ran at higher speeds on the first day of wheel access than females initiating VWR outside of Pro. Remarkably, initiating VWR during Pro also produced persistent increases in running distance, duration, speed, and escalation across subsequent weeks, despite normal cycling through other estrous phases. In contrast, exogenous estradiol (E2) administered at VWR initiation did not alter day-1 behavior, but increased running distance and duration across subsequent weeks without affecting running speed or escalation. To determine whether dorsal striatal dopamine contributes to these effects, we inhibited the substantia nigra (SN) to dorsolateral striatum (DLS) pathway on the first day of VWR. This manipulation reduced the immediate and long-term effects of initiating VWR during Pro on running duration and distance but not speed or escalation. These findings identify behavioral initiation as a critical window during which hormones and nigrostriatal signaling influence future engagement in physical activity. Furthermore, analysis of individual components of VWR architecture reveals that distinct features of VWR behavior can be dissociated mechanistically and thus could be used to investigate separate motivational processes underlying physical activity.

## Introduction

Physical inactivity is a major risk factor for premature mortality and chronic disease [1], yet a large proportion of adults fail to meet minimum physical activity recommendations [2]. Females consistently exhibit higher rates of physical inactivity than males across regions and age groups [2–5]. Developing stable exercise habits may help prevent and mitigate the adverse effects of a sedentary lifestyle [6–9], as individuals with stronger exercise habits are more likely to maintain long-term physical activity [8, 10–12]. Identifying mechanisms that promote exercise habit formation could therefore have substantial public health impact.

Voluntary wheel running (VWR) in rodents provides a powerful model for studying the development of intrinsic exercise habits. When given wheel access, rodents exhibit a characteristic pattern of behavior in which running distances progressively increase (escalation phase) before stabilizing at high levels (maintenance phase). This escalation-to-maintenance progression parallels the transition from goal-directed to habitual control of appetitive operant behavior, which is associated with a shift from dependence on the nucleus accumbens (NAc) and dorsomedial striatum (DMS) to engagement of the dorsolateral striatum (DLS) [13, 14]. Consistent with this framework, VWR depends on the DMS during escalation but shifts to DLS dependence during maintenance [15]. Female rodents run farther, longer, and faster than males [15–17], escalate to the maintenance phase more rapidly [15, 18], and rely on the DLS to govern VWR earlier after the start of VWR compared to males [15]. Together, these findings suggest that sex differences in physical activity may reflect sex-specific mechanisms of habit formation, positioning VWR as a useful model for identifying factors that promote persistent exercise behavior [14].

VWR in females is strongly influenced by ovarian hormones, which fluctuate across the estrous cycle. Estradiol (E2) levels peak during proestrus (Pro) and are lowest during metestrus [19]. Females in Pro run greater distances than females in other estrous phases [15, 16, 18, 20, 21], whereas ovariectomy abolishes cyclical running patterns and reduces running behavior and escalation rates to male-like levels [18]. E2 appears to mediate these effects, as E2 replacement restores typical female VWR behavior in ovariectomized animals [17, 22].

Despite these findings, important questions remain unresolved. Because prior studies have focused primarily on cumulative running measures, it remains unclear whether ovarian hormones influence the acquisition or expression of VWR. It is also unknown whether fluctuations in E2 across days are required for rapid escalation, or whether elevated E2 at the onset of wheel access is sufficient to promote long-term female-typical running patterns. Elevated E2 facilitates learning, dopamine (DA)-dependent reinforcement, and memory consolidation [23, 24], raising the possibility that VWR initiated during high-hormone states is more readily encoded and consolidated into habitual behavior. A likely mediator of this effect is substantia nigra (SN) DA input to the DLS, as Pro and E2 enhance DLS DA release [25–27] and plasticity [28], and medium-spiny neurons in the DLS expressing D1 receptors (D1-MSNs), which are low-affinity DA receptors sensitive to high concentrations of DA [29], play a key role in habit formation [30].

In the present study, we examined how estrous phase at exercise initiation influences both immediate and long-term VWR behavior. We quantified multiple features of running architecture, including distance, duration, speed, and bout number, which may reflect distinct motivational processes. Females initiating VWR during Pro displayed greater running on the first day of wheel access and sustained increases in running behavior across subsequent weeks. Pro initiation was also associated with a shift in D1-MSN activity toward the DLS relative to the DMS. Finally, the effects of Pro on specific aspects of running architecture were recapitulated by E2 administration and prevented by chemogenetic inhibition of the SN–DLS DA pathway. Together, these findings reveal a mechanism through which ovarian hormones promote the development of persistent exercise behavior in females.

## Methods

### Animals

Adult male and female Long Evans rats (Charles River, Wilmington, MA) were single-housed in Plexiglas cages (45.5 × 24 × 21 cm) with locked running wheels (Starr Life Sciences, Oakmont, PA). Animals were maintained on a 12 h light-dark cycle (0600–1800), at 22°C and 30% humidity, with ad libitum access to food (Teklad 2020X; Envigo) and water. Rats acclimated for 1 week before experimentation. All procedures were approved by the University of Colorado Denver Institutional Animal Care and Use Committee.

### Voluntary Wheel Running

Wheels in the cages of each rat in VWR groups were unlocked at the start of the active cycle on day 1 and remained unlocked henceforth. VWR was recorded automatically every 1 min using VitalView software (Starr Life Sciences). Distance was calculated by multiplying revolutions by wheel circumference and other features of VWR architecture (time spent running, running speed, and number of running bouts) were calculated with excel macros [18]. A running bout was defined as ≥3 revolutions within a 1-min period.

### Vaginal Lavage and Cell Cytology

Estrous cycle phase was determined by daily vaginal lavage and cell morphology for 8 days prior to the onset of VWR and again on the first day of VWR, 1 hour before wheel unlocking, as previously described [15, 25]. Daily lavage prior to VWR enabled prediction of estrous phase on day 1, facilitating group assignment and experimental planning. Rats were not started on VWR during estrus because E2 levels are intermediate during this phase.

### Fluorescent In Situ Hybridization

Female rats either in Pro or Not Pro on day 1 of VWR ran for 3 d and then, on the 4^th^ d of VWR, were euthanized ∼20 min after the start of the active cycle by rapid decapitation. Data from this experiment have been published previously [15], but here, we analyzed the previously published data based on the phase during which females started VWR. As such, protocols for fluorescent *in situ* hybridization for *drd1* and *cfos* mRNAs, imaging, and quantification can be found in our prior publication [15].

### Surgical Procedures

All surgeries were performed under ketamine (75 mg/kg, i.p.) and medetomidine (0.5 mg/kg, i.p.) anesthesia. Carprofen (5 mg/kg, s.c.) and penicillin G (22,000 IU/rat, s.c.) were administered at induction and every 24 h for 72 h postoperatively. Rats recovered for at least 2 weeks before experimentation. Viral infusions were delivered bilaterally using Hamilton syringes at 0.1 µL/min. For chemogenetic studies, all rats received AAV2/retro-eSYN-EGFP-T2A-iCre-WPRE (Vector Biolabs, Cat# VB4855; 1 µL/side) into the DLS (+0.5 mm AP, ±3.9 mm ML, −5.4 mm DV) and either pAAV8-hSyn-DIO-mCherry (Addgene #50459; http://n2t.net/addgene:50459; 1 µL/side) or pAAV8-hSyn-DIO-hM4Di(Gi)-mCherry (Addgene #44362; http://n2t.net/addgene:44362; 1 µL/side) into the SN (−5.4 mm AP, ±3.0 mm ML, −8.4 mm DV). mCherry expression was amplified by immunohistochemistry as previously described [25], and viral expression in the DLS and SN was verified in all rats. Animals with missed injections or insufficient viral expression were excluded. This intersectional viral strategy suppresses electrically evoked DLS dopamine release by ∼60% [25].

### Drugs

Rats received saline (1 mL/kg, i.p.) or JHU37160 dihydrochloride (J60; Hello Bio, Cat# HB6261; 0.1 mg/kg, i.p.) 30 min before the active cycle on the first day of VWR. E2 (Sigma Aldrich, Cat# E8875-1G; 4.5 µg/0.1 mL in sesame oil vehicle, Cat# S3547) or vehicle was administered as a single intrascapular s.c. injection (0.1 mL/kg) 30 min before wheel access. Both solutions were prepared immediately before use.

### Serum Estradiol Measurement

Trunk blood was collected in EDTA tubes from cycling female rats either during Metestrus (n = 13), Pro (n = 5), or 30 min following E2 administration (n = 9). Samples were spun in a centrifuge for 15 min (3000 g, 4°C) and supernatant was collected and stored at -80°C. E2 was measured using ELISA (ALPCO, Cat# 55-ESTRT-E01) according to the manufacturer’s instructions.

### Data Analysis

The effects of treatments were analyzed with repeated measures (10-min, hourly, and weekly running data), one-way (E2, average running data), or two-way (average running data in Figures 3 and 4) ANOVA, as appropriate, using GraphPad Prism (x.X). Main effects and interactions were considered significant if *p* < 0.05. When appropriate, post-hoc analyses were performed with Tukey’s multiple comparison tests. BioRender was used in the creation of Figures 3 and 5.

## Results

### Estrous phase at initiation impacts VWR architecture on the first day of wheel access and increases subsequent VWR

To determine if estrous phase variations in VWR behavior emerge over time or are present upon initial wheel exposure, and whether estrous phase at VWR initiation influences subsequent behavior, we compared VWR architecture during the first active cycle of wheel access and across subsequent weeks between rats starting VWR in Pro or Not Pro. Consistent with previous studies [15, 16, 18, 20, 21], rats in metestrus and diestrus did not differ on day 1 and were therefore combined into a single Not Pro group (n = 13) for comparison with rats in Pro (n = 11).

Rats initiating VWR in Pro ran greater distances during the first active cycle (F(1,22)=6.1; p=0.02; Figure 1A), an effect that emerged within the first hour of wheel access (main effect of phase: F(1,22)=4.05; p=0.05; Figure 1B). Increased distance was driven by greater time spent running (F(1,22)=4.7; p=0.04; Figure 1C) and higher running speed (F(1,22)=4.83; p=0.03; Figure 1E). Differences in running duration were evident during the first hour (main effect of phase: F(1,22)=5.07; p=0.03; Figure 1D), whereas running speed was not (main effect of phase: F(1,22)=0.44; p=0.52; Figure 1F). The number of running bouts did not differ between groups during either the full active cycle (F(1,22)=1.43; p=0.24; Figure 1G) or the first hour (main effect of phase: F(1,22)=0.04; p=0.85; Figure 1H). Across subsequent weeks, distance run (F(2,44)=77.59; p<0.0001; Figure 1E), time spent running (F(2,44)=42.78; p<0.0001; Figure 1F), and running speed (F(2,44)=158.8; p<0.0001; Figure 1G) increased in both groups, whereas running bouts declined (F(2,44)=48.61; p<0.0001) at a similar rate (phase × time: F(2,44)=1.6; p=0.21; Figure 1H). Rats that began VWR in Pro ran farther (F(1,22)=7.73; p=0.01) and showed greater escalation of running distance (phase × time: F(2,44)=3.34; p=0.04; Figure 1E). This effect was attributable to greater running duration (F(1,22)=6.5; p=0.01; Figure 1F) and speed (F(1,22)=5.9; p=0.02; Figure 1G), but not running bouts (F(1,22)=0.79; p=0.38; Figure 1H). Running speed escalated more rapidly in Pro animals (phase × time: F(2,44)=3.7; p=0.03), whereas running duration did not (phase × time: F(2,44)=0.6; p=0.55). Estrous phase at VWR initiation did not affect inactive-cycle activity (Supplemental Figure 1) or circadian rhythmicity (Supplemental Figure 2). Together, these findings indicate that estrous phase at VWR onset influences both the initial expression and long-term escalation of running distance, duration, and speed.

**Figure 1.**
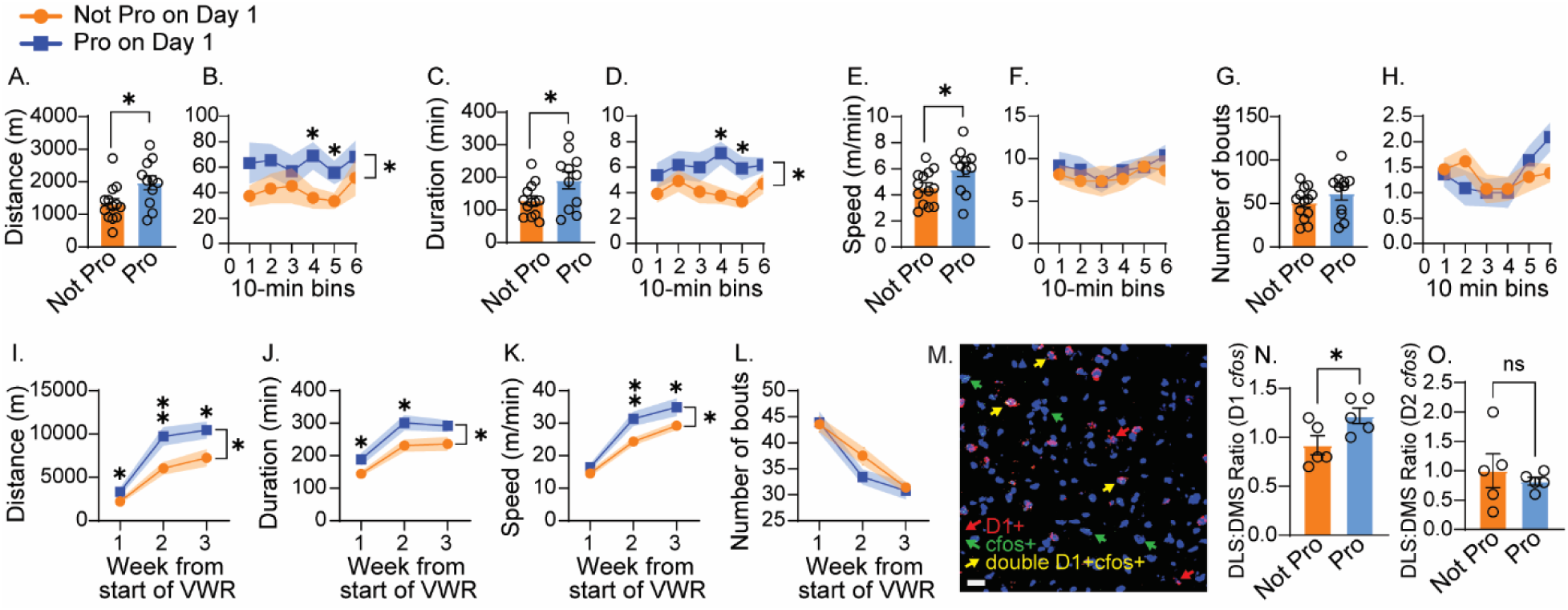
Estrous cycle phase at the onset of voluntary wheel running (VWR) influences running architecture on day 1 and across subsequent weeks. Cycling female rats housed with locked running wheels initiated VWR at the start of the dark (active) cycle while in either proestrus (Pro) or metestrus/diestrus (Not Pro). (A) Distance run during the active cycle on day 1. (B) Distance run during the first hour in 10-min bins. (C) Time spent running during the active cycle on day 1. (D) Time spent running during the first hour in 10-min bins. (E) Average running speed during the active cycle on day 1. (F) Running speed over the first hour in 10-min bins. (G) Number of running bouts during the active cycle on day 1. (H) Number of running bouts in the first hour in 10-min bins. Across subsequent weeks, running behavior is expressed as average daily values during the active cycle for distance (I), time spent running (J), speed (K), and number of bouts (L). (M) Representative image of FISH labeling. (N-O) Dorsolateral striatum (DLS): dorsomedial striatum (DMS) expression ratios for D1- (N) and single *cfos* (putative D2) - expressing (O) cells by estrous phase on day 1. Asterisks denote significant post hoc comparisons (*p < 0.05, **p < 0.01; ns, not significant). Data are presented as mean ± SEM, with open circles indicating individual data points and shaded lines reflecting SEM.

We previously reported that a bout of VWR increases *cfos* mRNA expression in D1-MSNs of the DLS, DMS, and NAc shell and that females show earlier recruitment of DLS D1-MSNs than males during VWR acquisition [15]. Reanalysis of those data by estrous phase at VWR initiation revealed that females starting VWR in Pro exhibited a higher DLS:DMS ratio of exercise-induced D1-MSN activity on day 4 than females starting in other phases (F(1,8)=5.69; p=0.04; Figure 1J). This effect was specific to D1-MSNs, as no difference was observed in cells expressing *cfos* alone (putative D2-MSNs; F(1,8)=0.37; p=0.55; Figure 1K). Initiating VWR in Pro also did not affect exercise-induced *cfos* expression in D1-MSNs (F(1,8)=0.81; p=0.39) or putative D2-MSNs (F(1,8)=4.06; p=0.08) in the NAc shell (data not shown). Together, these findings indicate that initiating VWR in Pro biases subsequent exercise-induced D1-MSN activity within the dorsal striatum toward the DLS.

### E2 administration on the first day of wheel access alters subsequent VWR architecture without impacting VWR on day 1

To examine the role of E2 in the effects of estrous phase at VWR initiation, cycling female rats received Vehicle (n=8) or E2 (4.5 µg/kg; n=8) 30 min before the start of the active cycle on the first day of wheel access. Daily vaginal lavage was used to minimize the number of rats in Pro on day 1 and these were distributed across groups; consequently, 2 Vehicle-treated and 3 E2-treated rats were in Pro at VWR onset. Because estrous phase at VWR initiation influences behavior for at least 3 weeks (Figure 1E), VWR was extended to 6 weeks to assess the persistence of E2 effects.

Relative to Vehicle, E2 administration did not affect distance run during the first active cycle (F(1,14)=0.25; p=0.62; Figure 2A) or first hour of wheel access (F(1,14)=0.12; p=0.74; Figure 2B), time spent running during the first active cycle (F(1,14)=0.02; p=0.88; Figure 2C) or first hour (F(1,14)=0.17; p=0.68; Figure 2D), running speed during the first active cycle (F(1,14)=2.62; p=0.12; Figure 2E) or first hour (F(1,14)=0.002; p=0.97; Figure 2F), or the number of running bouts during the first active cycle (F(1,14)=0.14; p=0.71; Figure 2G) or first hour (F(1,14)=0.02; p=0.89; Figure 2H). Despite the lack of acute effects, E2 altered subsequent VWR behavior. Across 6 weeks, running distance (F(5,70)=48.05; p<0.0001; Figure 2E), time spent running (F(5,70)=36.5; p<0.0001; Figure 2F), and running speed (F(5,70)=84.77; p<0.0001; Figure 2G) increased, whereas running bouts decreased (F(5,70)=4.72; p=0.0009; Figure 2H), regardless of group. Compared with Vehicle, E2-treated rats ran farther (F(1,14)=4.77; p=0.04; Figure 2E) and spent more time running (F(1,14)=6.11; p=0.02; Figure 2F), but did not differ in running speed (F(1,14)=0.23; p=0.63; Figure 2G) or bout number (F(1,14)=0.0001; p=0.99; Figure 2H). No drug × time interactions were detected (distance: F(5,70)=0.26; p=0.93; duration: F(5,70)=0.79; p=0.56; speed: F(5,70)=0.63; p=0.67; bouts: F(5,70)=0.14; p=0.98), indicating that E2 at initiation did not facilitate escalation. E2 also had no effect on inactive-cycle activity (Supplemental Figure 3) or circadian rhythmicity (Supplemental Figure 4). Circulating E2 levels differed between groups (F(2,24)=3.62; p=0.04), with the highest levels observed following E2 administration (Figure 2M). Together, these findings indicate that E2 administration at VWR initiation does not acutely alter running behavior but produces a delayed increase in running distance and duration without affecting running speed or escalation.

**Figure 2.**
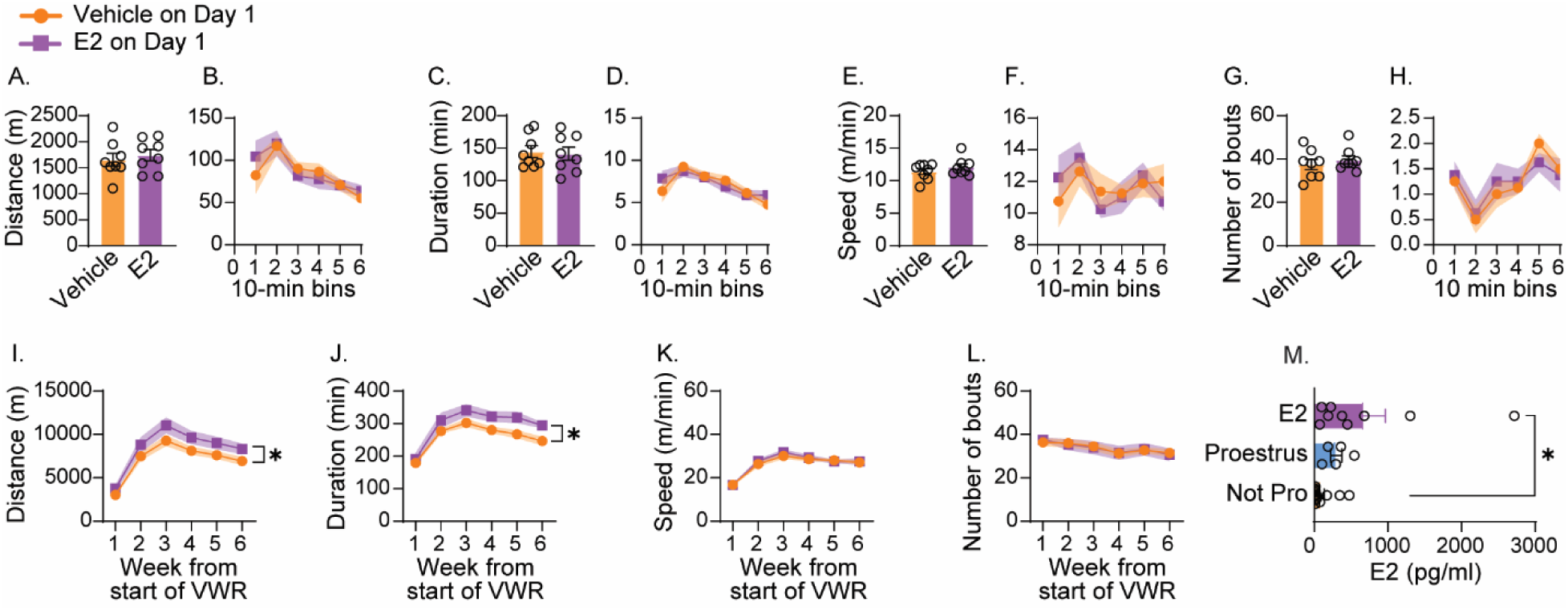
Effects of estradiol (E2) on voluntary wheel running architecture on the first day of wheel access and across subsequent weeks. Freely cycling female rats housed with locked running wheels received vehicle or E2 (4.5 µg/kg) 30 min prior to the onset of the active (dark) cycle, at which point wheels were unlocked to allow voluntary running. (A) Distance run during the active cycle on day 1. (B) Distance run during the first hour in 10-min bins. (C) Time spent running during the active cycle on day 1. (D) Time spent running during the first hour in 10-min bins. (E) Average running speed during the active cycle on day 1. (F) Running speed over the first hour in 10-min bins. (G) Number of running bouts during the active cycle on day 1. (H) Number of running bouts in the first hour in 10-min bins. Across subsequent weeks, running behavior is expressed as average daily values during the active cycle for distance (I), time spent running (J), speed (K), and number of bouts (L). (M) Plasma E2 levels measured from trunk blood collected during metestrus/diestrus (Not Pro) or proestrus (Pro), or 30 min following exogenous E2 administration, confirming differences in circulating hormone levels across conditions. Asterisks denote significant group differences (*p < 0.05). Data are presented as mean ± SEM, with open circles indicating individual data points and shaded lines reflecting SEM.

### The long-lasting effect of initiating VWR in Pro on VWR architecture is dependent on SN-DLS pathway activity on day 1

Because elevated E2 enhances stimulus-evoked DA release in the DLS [25, 27, 31] and DLS DA signaling contributes to habit formation [32, 33], we hypothesized that SN-DLS pathway activity mediates the effects of Pro on both initial and subsequent VWR behavior. To test this, we used an intersectional chemogenetic approach to inhibit SN neurons projecting to the DLS on the first day of VWR in females initiating running in either Pro or Not Pro (Figure 3A). Viral expression was robust in both the SN and DLS (Figures 3B,C). After excluding four animals with missed viral injections and one in pseudopregnancy, final group sizes were: Not Pro/mCherry (n=11), Not Pro/GiDREADD (n=7), Pro/mCherry (n=10), and Pro/GiDREADD (n=9).

**Figure 3.**
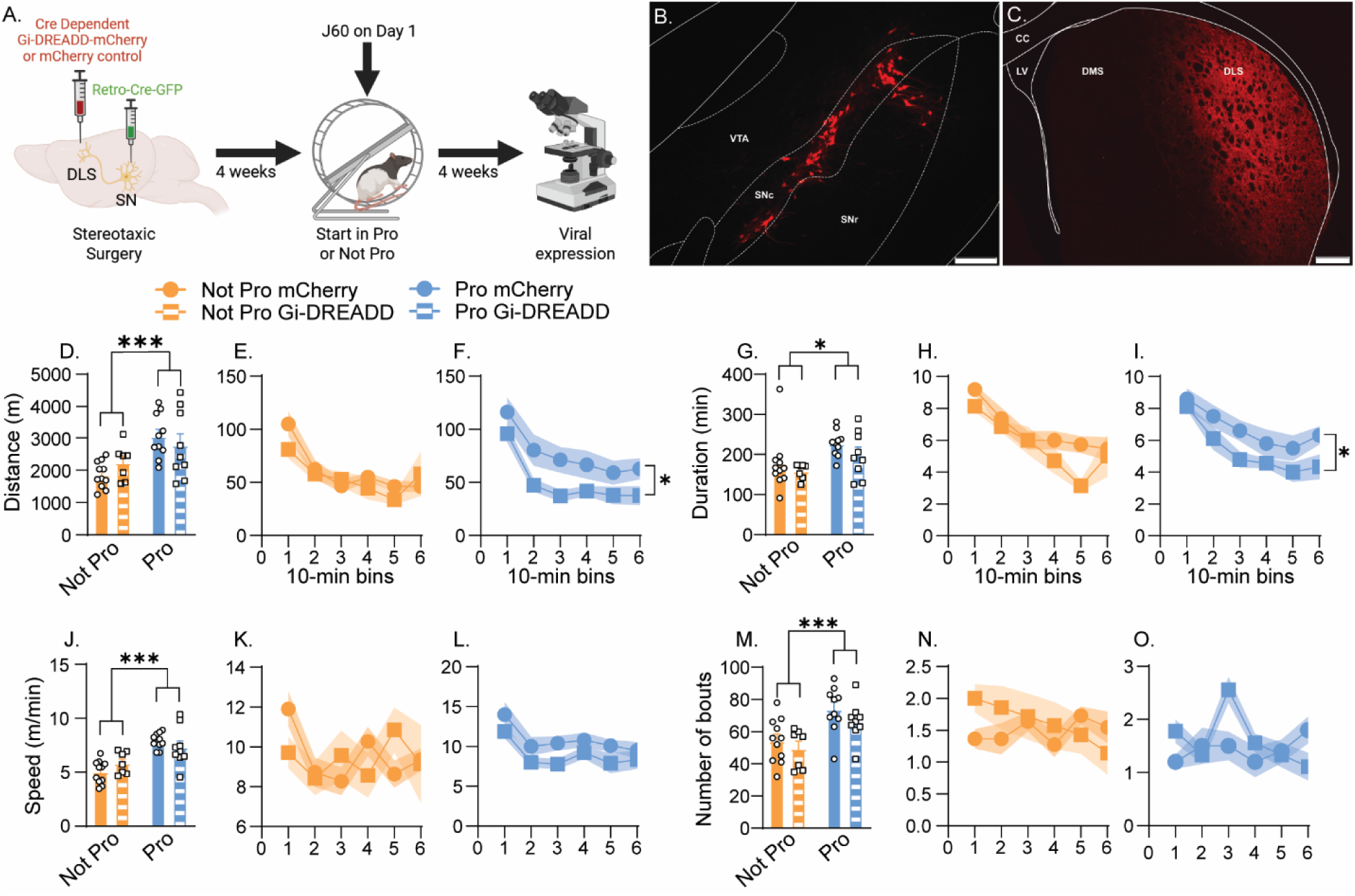
Effects of chemogenetic inhibition of the substantia nigra (SN) – dorsolateral striatum (DLS) pathway on voluntary wheel running (VWR) architecture on the first day of wheel access. (A) Experimental design. Viral mCherry expression in the SN (B) and DLS (C). Average distance run during the first active cycle (D) or during the first hour in rats initiating VWR in Not Pro (E) or Pro (F). Average time spent running during the first active cycle (G) or during the first hour in rats initiating VWR in Not Pro (H) or Pro (I). Average running speed during the first active cycle (J) or during the first hour in rats initiating VWR in Not Pro (K) or Pro (L). Average number of running bouts during the first active cycle (M) or during the first hour in rats initiating VWR in Not Pro (N) or Pro (O). Asterisks denote significant main effects of day 1 phase or viral expression (ANOVA; *p < 0.05; ***p<0.001). Data are presented as mean ± SEM, with circles or squares indicating individual data points and shaded lines reflecting SEM.

Rats initiating VWR in Pro ran farther (F(1,33)=13.31; p=0.0009; Figure 3D), spent more time running (F(1,33)=7.47; p=0.01; Figure 3G), ran faster (F(1,33)=29.24; p<0.0001; Figure 3J), and completed more bouts (F(1,33)=16.79; p=0.0003; Figure 3M) during the first active cycle than rats initiating VWR in Not Pro. SN-DLS inhibition did not affect distance (F(1,33)=0.04; p=0.84; Figure 3D), duration (F(1,33)=1.86; p=0.18; Figure 3G), speed (F(1,33)=0.08; p=0.77; Figure 3J), or bouts (F(1,33)=2.98; p=0.09; Figure 3M) across the full active cycle (all phase × virus interactions p>0.05). However, analysis of the first hour in 10-min bins revealed that SN-DLS inhibition reduced running distance (F(1,33)=4.43; p=0.04; Figures 3E,F) and duration (F(1,33)=6.99; p=0.01; Figures 3H,I). These effects were restricted to rats initiating VWR in Pro, in which inhibition reduced distance (F(1,17)=4.90; p=0.04; Figure 3F) and duration (F(1,17)=4.95; p=0.03; Figure 3I). In contrast, inhibition had no effect on distance (F(1,16)=0.37; p=0.55; Figure 3E) or duration (F(1,16)=2.32; p=0.15; Figure 3H) in rats initiating VWR in Not Pro. Neither running speed (F(1,33)=2.05; p=0.16; Figures 3K,L) nor bout number (F(1,33)=0.15; p=0.70; Figures 3N,O) was affected by SN-DLS inhibition during the first hour, regardless of phase. Together, these findings indicate that SN-DLS inhibition transiently reduces running distance and duration in Pro females during the first hour of VWR without affecting overall day 1 activity.

Rats initiating VWR in Pro continued to exhibit greater running over subsequent weeks, covering greater distances (F(1,33)=16.23; p=0.0003; Figure 4A–C), spending more time running (F(1,33)=9.46; p=0.004; Figure 4D–F), and running at higher speeds (F(1,33)=10.31; p=0.003; Figure 4G–I), with no difference in bout number (F(1,33)=0.19; p=0.67; Figure 4J–L). This persistent effect depended on SN–DLS activity on day 1, as indicated by a significant phase × virus interaction for distance (F(1,33)=4.53; p=0.04) and similar trends for duration (F(1,33)=3.20; p=0.08) and speed (F(1,33)=3.04; p=0.09). Distance (phase × time: F(3,99)=7.64; p=0.0001), duration (phase × time: F(3,99)=7.11; p=0.0002), and speed (phase × time: F(3,99)=6.15; p=0.0007) also escalated more rapidly in rats initiating VWR in Pro, whereas SN–DLS inhibition did not alter escalation rates (all virus × time interactions p>0.05). No main effect of virus was detected for any VWR measure (all p>0.05), and neither estrous phase nor SN–DLS inhibition affected inactive-cycle activity (Supplemental Figure 5). Together, these findings indicate that SN–DLS activity during Pro at VWR initiation is required for the persistent increase in VWR, primarily through greater time spent running, but not for enhanced running speed or accelerated escalation.

**Figure 4.**
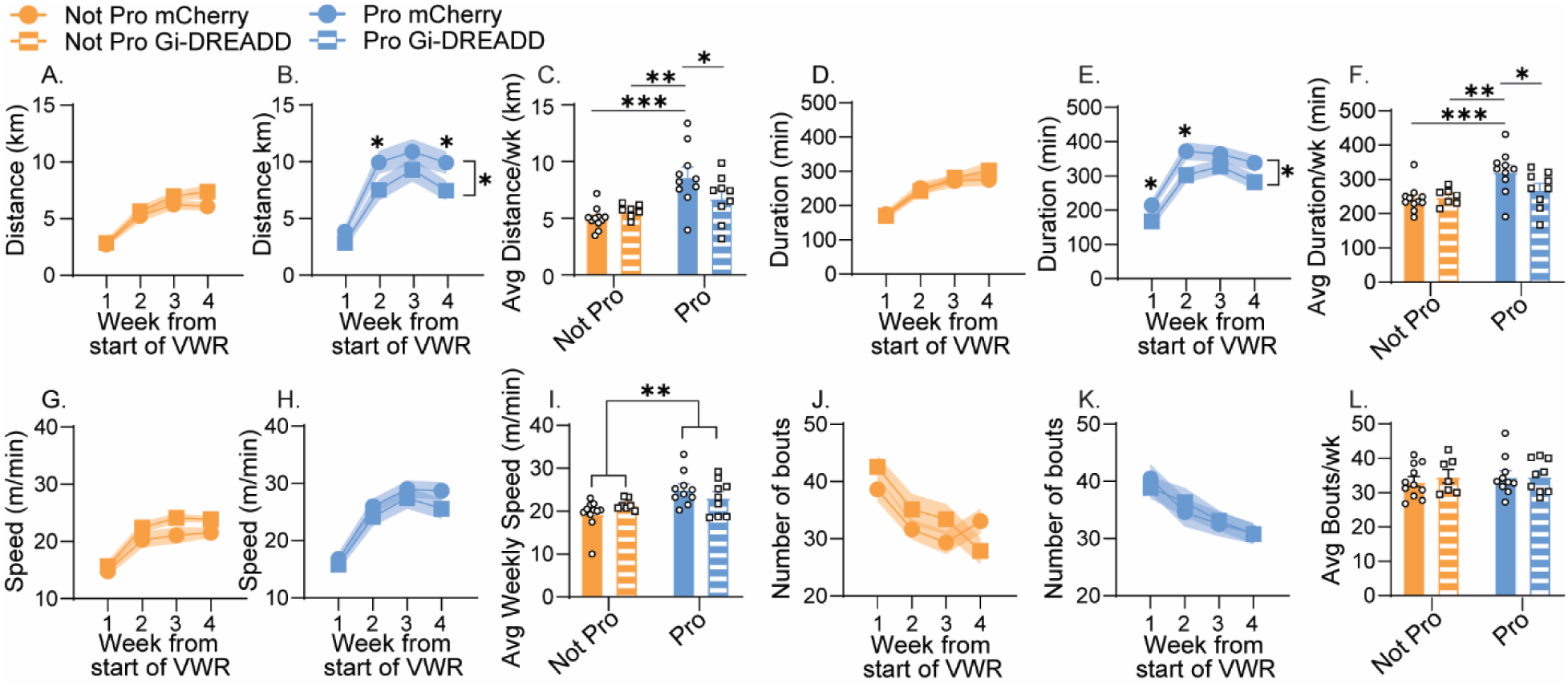
Effects of chemogenetic inhibition of the substantia nigra (SN) – dorsolateral striatum (DLS) pathway on voluntary wheel running (VWR) architecture across weeks. Experimental design is shown in Figure 3A. Distance run across weeks in rats initiating VWR in Not Pro (A) or Pro (B). Average weekly distance run (C). Time spent running across weeks in rats initiating VWR in Not Pro (D) or Pro (E). Average weekly distance run (F). Running speed across weeks in rats initiating VWR in Not Pro (G) or Pro (H). Average weekly running speed (I). Number of running bouts across weeks in rats initiating VWR in Not Pro (J) or Pro (K). Average weekly running bouts (L). Asterisks denote significant main effects of day 1 estrous phase or virus (ANOVA), or Bonferroni post-hoc comparisons (*p < 0.05; **p<0.01 ***p<0.001). Data are presented as mean ± SEM, with circles or squares indicating individual data points and shaded lines reflecting SEM.

## Discussion

Previous studies have reported estrous cycle-dependent differences in VWR, but it remains unclear whether these effects emerge immediately or require prior experience, and whether hormone state at exercise initiation influences long-term VWR. Because E2 potentiates DA release in the DLS [25–27], and females are more prone than males to habit formation [34, 35] and DLS-dependent control of VWR [15], we hypothesized that initiating VWR during Pro would enhance both immediate and long-term running behavior in female rats. Consistent with this hypothesis, estrous-related differences in VWR were evident on the first day of wheel access, indicating they do not require prior running experience. Moreover, initiating VWR under high ovarian hormone conditions increased subsequent running behavior, an effect that was partially dependent on SN–DLS pathway activity.

We analyzed components of VWR because they may reflect distinct motivational processes. Time spent running may indicate motivation to sustain activity, whereas running speed may reflect vigor, or energetic investment in performance [36, 37]. The number of running bouts may capture fragmentation of running behavior, although its motivational significance is less clear. Since estrous phase effects were driven by increases in running duration and speed, but not running bouts, this framework suggests that the high ovarian hormone state characteristic of Pro increases motivation for sustained activity and vigor without altering patterning. The rapid emergence of these effects after wheel access further suggests that ovarian hormone state influences baseline motivation for physical activity, rather than only learned or habitual VWR behavior. Interestingly, however, female rats in Pro do not express greater locomotor activity while exploring a novel environment than rats in Not Pro [25], suggesting that the acute effect of Pro could be specific to motivation to engage in VWR, rather than non-selective enhancements in locomotor performance, *per se*. Consistent with this, stimulation of the SN-DLS pathway also does not increase locomotor activity in a novel environment [25].

Because exercise behavior during the adoption phase predicts long-term maintenance and adherence [38, 39], initiating physical activity during high-hormone states may create more reinforcing early experiences that promote continued engagement. Consistent with this idea, hormonal state at first wheel exposure influenced not only immediate VWR but also subsequent behavior. Females that began VWR in Pro increased running more rapidly and exhibited greater running duration, speed, and distance over subsequent weeks, despite cycling through other estrous phases. These effects were limited to the active phase and did not alter circadian rhythms, suggesting that Pro amplifies normal activity patterns rather than disrupting them. Together, these findings indicate that ovarian hormones at activity initiation influence the subsequent development of habitual physical activity, not merely its acute expression. Supporting this interpretation, females that initiated VWR during Pro showed a bias toward exercise-induced recruitment of D1-MSNs in the DLS, a neural substrate of habitual behavior [30, 32, 33].

Unlike Pro, elevating E2 at VWR initiation did not reproduce the day 1 increase in running, enhance running speed, or accelerate the escalation of VWR across weeks. These findings suggest that E2 alone is insufficient to drive the immediate increases in VWR associated with Pro. This may reflect the absence of coordinated E2-progesterone signaling characteristic of Pro, as progesterone enhances striatal DA release in the presence of E2 [31, 40, 41] and interacts with E2 to regulate MSN physiology [42]. Alternatively, exogenous E2 may engage signaling pathways distinct from those activated by the physiological hormone fluctuations that occur during Pro [28].

Despite lacking acute effects, E2 increased time spent running over subsequent weeks, resulting in greater overall distance run and suggesting a delayed influence on motivational systems underlying VWR. This delayed effect may reflect gradual changes in DA receptor sensitivity or expression [31, 43, 44]. Together, these findings suggest that Pro and E2 enhance VWR through distinct mechanisms: Pro promotes immediate, high-vigor performance and accelerates the development of vigorous activity, whereas E2 gradually increases overall engagement. More broadly, these results indicate that exercise duration, vigor, and escalation are dissociable motivational processes that can be differentially influenced by hormonal state at activity onset.

These findings prompted us to test whether activity of the SN-DLS pathway, the primary source of DA input to the DLS, at the time of initial VWR exposure contributes to the acute effects of Pro, their persistence, or both. We focused on the DLS because it is critical for habit formation [30, 33] and because initiating VWR during Pro biases subsequent D1-MSN recruitment toward this region. However, because some SN neurons release neurotransmitters other than DA [45] and project to both the DLS and DMS, the results should be interpreted with this limitation in mind.

Contrary to our expectations, SN-DLS inhibition did not alter VWR architecture during the first active cycle, possibly because J60 produced maximal inhibition only during the first few hours after administration [46]. Consistent with this interpretation, SN-DLS inhibition reduced running distance and duration during the first hour of VWR in Pro, but not Not Pro, animals. These findings suggest that SN-DLS activity is required for the acute increase in motivation to sustain VWR associated with Pro, while playing little role in initial VWR acquisition under low-hormone conditions.

Notably, SN-DLS inhibition did not affect running speed either acutely or across subsequent weeks. Instead, its effects on both short- and long-term VWR were driven by reductions in time spent running, leading to lower running distances in animals that initiated VWR during Pro. This dissociation is consistent with evidence that the DLS supports action selection and behavioral persistence [32, 47], whereas movement vigor depends on broader DA-related basal ganglia circuits [36, 48]. Together, these findings suggest that SN-DLS activity at VWR initiation mediates the effects of Pro on motivation to sustain physical activity and contributes to the long-term behavioral consequences of this hormonal state.

It is particularly interesting that both E2 administration and SN-DLS inhibition affected running distance selectively through running duration and not speed. This pattern suggests that the effects of Pro on VWR behavior may be mediated by multiple neurobiological mechanisms, summarized in Figure 5. One possibility is that elevated E2 during Pro enhances SN-DLS signaling, thereby increasing the motivation to sustain VWR, which is reflected in greater running duration. In contrast, the increased running speed and accelerated escalation observed in rats starting VWR in Pro may be mediated by other hormonal or neural mechanisms, such as DA signaling in other striatal regions, that are not reproduced by E2 administration alone and are independent of SN-DLS signaling. These findings further support the idea that distinct components of VWR architecture reflect dissociable motivational processes that can be regulated independently. Future research is needed to clarify the mechanisms underlying the enhanced vigor and escalation seen in rats initiating VWR in Pro.

**Figure 5.**
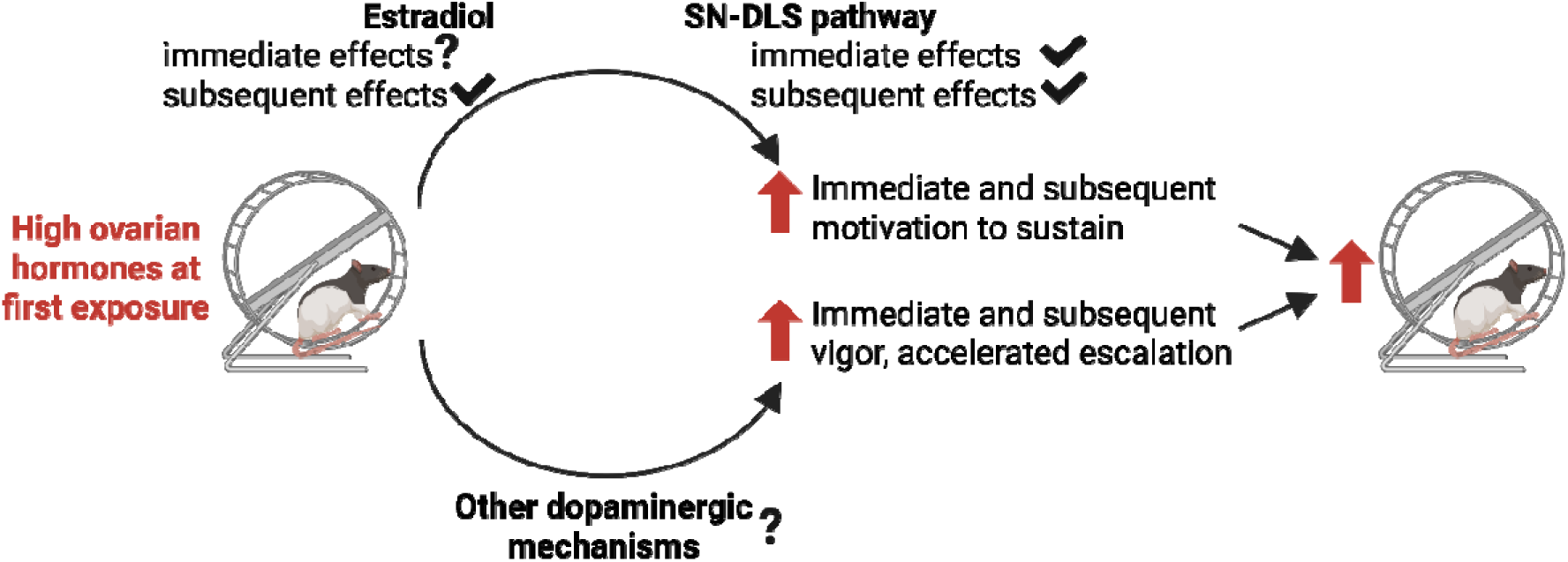
Schematic of the role of estradiol and the substantia nigra (SN) to dorsolateral striatum (DLS) pathway in mediating the effects of high ovarian hormones during the first exposure to running wheels on immediate and subsequent voluntary exercise.

The current results may have important implications for understanding physical activity in women across the lifespan. High-ovarian hormone states may provide a biological context that enhances both the initial experience of exercise and its long-term persistence. Conversely, conditions associated with reduced ovarian hormone fluctuations, such as menopause, are associated with alterations in physical activity patterns and exercise behavior in women [49]. The present findings raise the possibility that one mechanism contributing to these effects is a reduction in opportunities for highly reinforcing initial exercise experiences, although this hypothesis remains to be tested directly.

## Conclusion

The present findings demonstrate that hormonal state at the onset of physical activity influences both immediate behavior and long-term engagement. These effects are partially mediated by E2 and interactions of ovarian hormones with the SN-DLS pathway. Analysis of micro-architecture of VWR behavior reveals that motivational aspects underlying VWR can be dissociable at a mechanistic level. These results identify behavioral initiation as a critical period during which endocrine and circuit-level factors determine whether physical activity becomes persistent and self-sustaining, with potential implications for formation of habitual exercise in humans.

## Acknowledgments

Portions of this manuscript were edited with assistance from a large language model (ChatGPT, OpenAI). The authors generated, reviewed, edited, and take full responsibility for all content.

## Data Availability Statement

The datasets generated during and/or analyzed during the current study are available from the corresponding author on reasonable request.

## Author contributions

MKT designed experiments, and contributed to the acquisition, analysis, and interpretation of data, and drafting of manuscript; KMK designed experiments and contributed to the acquisition, analysis, and interpretation of data, and drafting of manuscript; AAH contributed to the acquisition of data; JRF contributed to the acquisition of data; JDW contributed to the acquisition and analysis of data; CSM contributed to the acquisition and analysis of data; BNG designed experiments and contributed to the analysis and interpretation of data, and drafting of manuscript. All authors contributed to critical revisions of the document, give their final approval for publication, and agree to be accountable for all aspects of the work in ensuring that questions related to the accuracy or integrity of any part of the work are appropriately investigated and resolved.

