## Supplemental Material for "Ovarian hormonal state at exercise initiation interacts with nigrostriatal circuitry to determine long-term voluntary exercise behavior"

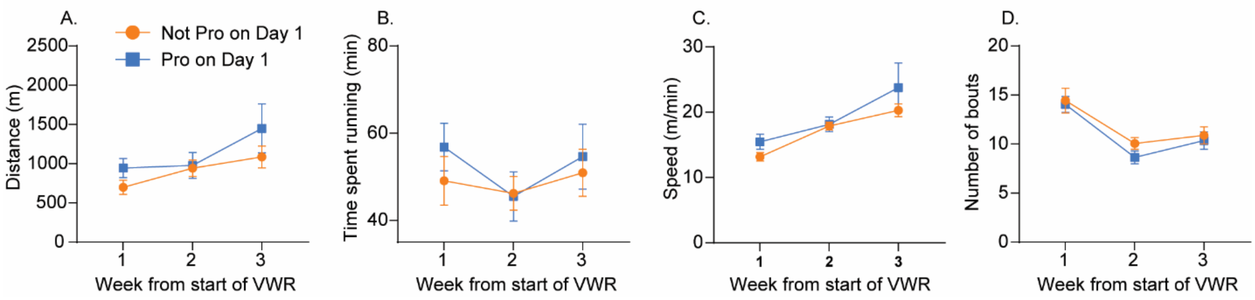


Supplemental Figure 1

During the inactive cycle, distance run ($F(2,44)=5.88$; $p=0.005$; A) and running speed ($F(2,44) = 13.4$; $p < 0.0001$; C), but not time spent running ($F(2,44)=1.9$; $p=0.16$; B) escalated over time. Similar to what was observed during the active cycle, number of running bouts during the inactive cycle decreased over time ($F(2,44)=20.817$; $p<0.0001$; D. However, unlike during the active cycle, the estrous phase during which VWR was initiated had no impact on VWR architecture during the inactive cycle. Neither the main effects of estrous phase (distance: $F(1,22)=1.49$; $p=0.23$; time spent running: $F(1,22)=3.22$; $p=0.57$; speed: $F(1,22)=0.05$; $p=0.81$; bout number: $F(1,22)=0.77$; $p=0.39$) nor the interactions between estrous phase and time (distance: $F(2,44)=0.77$; $p=0.46$; time spent running: $F(2,44)=0.52$; $p=0.59$; speed: $F(2,44)=1.66$; $p=0.21$; bout number: $F(2,44)=0.25$; $p=0.77$) differed significantly between groups during the inactive cycle. Data points are means ± SEM.

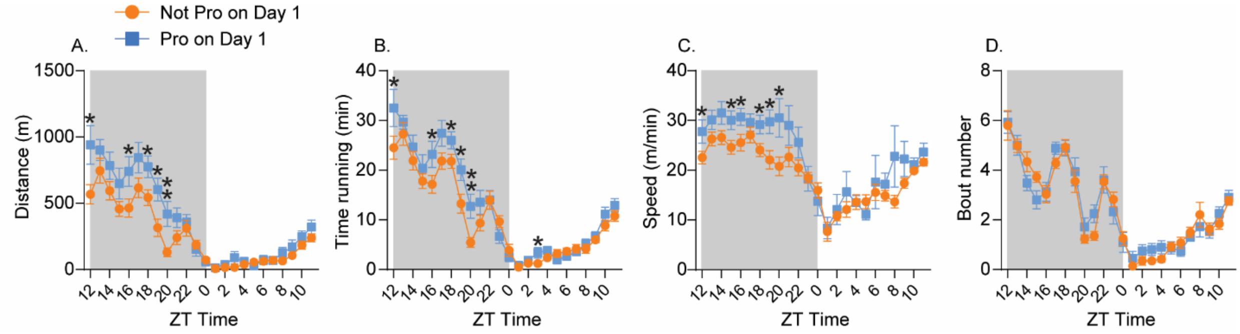


Supplemental Figure 2

In addition to running greater distances compared to rats in other phases, while in Pro, females have been reported to have altered circadian rhythmicity of voluntary wheel running (VWR). Specifically, females will begin running several hours prior to the start of the active cycle while in Pro ( ). To gain a more accurate picture of the impact of estrous phase on the first day of VWR on the hourly circadian rhythm of VWR over subsequent weeks, aspects of VWR architecture were analyzed per hour, collapsed across weeks, and compared between rats that started VWR in Pro and Not Pro.

Cycling female rats were given voluntary access to running wheels for 3 weeks. Rats were divided into those in the proestrus (Pro) or the metestrus or diestrus (Not Pro) phases of their estrous cycle on the first day of wheel access. Hourly running distance (A), time spent running (B), running speed (C), and bout number (D) were averaged across the 3 weeks. Distance ($F(23,506) = 57.92$; $p < 0.0001$; A), time spent running ($F(23,506) = 84.55$; $p < 0.0001$; B), running speed ($F(23,506) = 28.79$; $p < 0.0001$; C), and number of running bouts ($F(23,506) = 56.5$; $p < 0.0001$; D) all displayed typical circadian rhythms characterized by greater activity during the dark cycle. Repeated measures ANOVA revealed significant interactions between estrous phase and time of day for distance ($F(23,506) = 2.7$; $p < 0.0001$; A), time spent running ($F(23,506) = 2.16$; $p = 0.001$; B), and running speed ($F(23,506) = 1.82$; $p = 0.01$; C), but not number of bouts ($F(23,506) = 1.09$; $p = 0.34$; D). Post hoc analyses revealed that rats that began VWR in Pro ran greater distances, for a longer duration, and at faster speeds during the active cycle than rats that started in other phases. This confirms that estrous phase during which female rats begin VWR influences later VWR architecture during the active cycle but does not impact the circadian rhythmicity of VWR. ZT = zeitgeber time. Dark (active) phase highlighted in gray, and light (inactive) phase highlighted in white. Stars over individual time points indicate significant post-hoc tests. * p<0.05; **p<0.01. Data points represent group means +/- SEM.

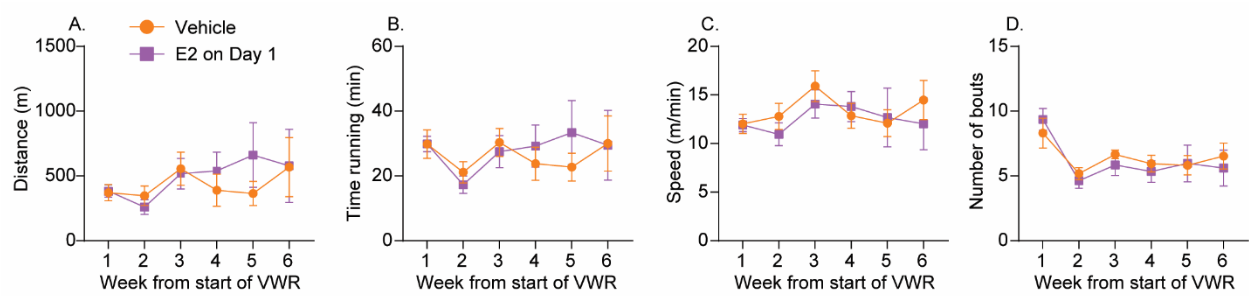


Supplemental Figure 3

Estradiol (E2) administration on the first day of voluntary wheel running (VWR) does not impact subsequent voluntary wheel running (VWR) behavior during the inactive cycle. Cycling female rats living with a locked running wheel were given Vehicle or E2 (xx) 30 min prior to the start of the active cycle. Wheels were unlocked at the start of the active cycle and rats were allowed to run voluntarily for 6 weeks. A) Average daily distance run during the inactive cycle. B) Average time spent running during the inactive cycle. C) Average running speed during the inactive cycle. D) Average number of running bouts during the inactive cycle. Neither distance run ($F(5,70) = 1.92$; $p = 0.1$; A), time spent running ($F(5,70) = 2.1$; $p = 0.07$; B), nor running speed ($F(5,70) = 1.91$; $p = 0.1$; C) during the inactive cycle significantly changed across weeks. Similar to what was observed during the active cycle, number of running bouts during the inactive cycle decreased over time ($F(5,70) = 9.7$; $p < 0.0001$; D). Similar to the lack of effect of estrous phase on initiation on subsequent VWR during the inactive cycle, E2 administration on the first day of VWR had no impact on VWR architecture during the inactive cycle. Neither the main effects of drug (distance: $F(1,14) = 0.11$; $p = 0.74$; time spent running: $F(1,14) = 0.04$; $p=0.83$; speed: $F(1,14) = 0.17$; $p = 0.68$; bout number: $F(1,14) = 0.07$; $p = 0.78$) nor the interactions between drug and time (distance: $F(5,70) = 0.91$; $p = 0.47$; time spent running: $F(5,70) = 0.98$; $p = 0.43$; speed: $F(5,70) = 0.72$; $p = 0.61$; bout number: $F(5,70) = 0.76$; $p = 0.57$) differed significantly between groups during the inactive cycle. Data points represent group means +/- SEM.

Supplemental Figure 4

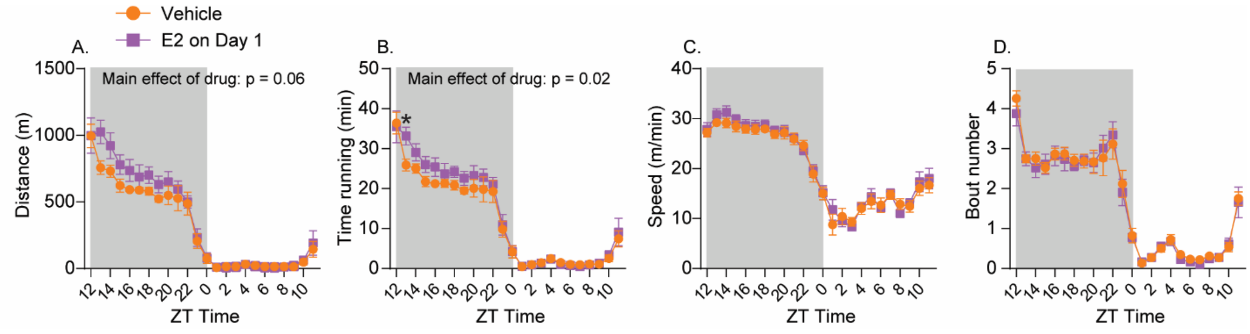


E2 administration on the first day of voluntary wheel running (VWR) alters subsequent VWR architecture during the active cycle but does not impact the circadian rhythmicity of VWR. Cycling female rats living with a locked running wheel were given Vehicle or E2 (xx) 30 min prior to the start of the active cycle. Wheels were unlocked at the start of the active cycle and rats were allowed to run voluntarily for 6 weeks. Hourly running distance (A), time spent running (B), running speed (C), and bout number (D) were averaged across the 6 weeks. Distance ($F(23,322) = 91.81$; $p < 0.0001$; A), time spent running ($F(23,322) = 101.72$; $p < 0.0001$; B), running speed ($F(23,322) = 108.11$; $p < 0.0001$; C), and number of running bouts ($F(23,322) = 92.64$; $p < 0.0001$; D) all displayed typical circadian rhythms characterized by greater activity during the dark cycle. E2 administration on day 1 of VWR increased hourly distance (although this effect just missed significance; $F(1,14) = 4.18$; $p = 0.06$; A) and time spent running ($F(1,14) = 6.29$; $p = 0.02$; B), but did not impact running speed ($F(1,14) = 0.31$; $p = 0.58$; C) or number of bouts ($F(1,14) = 0.15$; $p = 0.69$; D). Post hoc analyses revealed significant differences between E2 and Vehicle during the active cycle, but not inactive cycle. These data suggest that, like starting VWR in Pro, E2 administration on day 1 of VWR does not impact the circadian rhythmicity of VWR. ZT = zeitgeber time. Dark (active) phase highlighted in gray, and light (inactive) phase highlighted in white. Stars over individual time points indicate significant post-hoc tests. * $p<0.05$; ** $p<0.01$. Data points represent group means +/- SEM.

Supplemental Figure 5

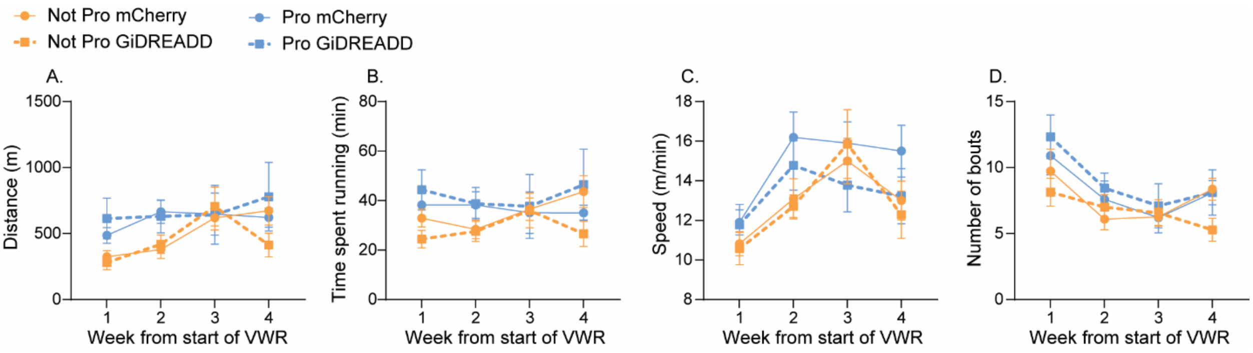


During the inactive cycle, distance run ($F(3,99) = 4.67$; $p = 0.004$; A) and running speed ($F(3,99) = 16.14$; $p < 0.0001$; C), but not time spent running ($F(3,99) = 0.67$; $p = 0.57$; B), escalated over time. Similar to what was observed during the active cycle, number of running bouts during the inactive cycle decreased over time ($F(3,99)=15.23$; $p<0.0001$; D. However, unlike during the active cycle, neither the estrous phase during which VWR was initiated (distance: $F(1,33) = 2.61$; $p = 0.11$; time spent running: $F(1,33)=1.57$; $p=0.22$; speed: $F(1,33) = 1.87$; $p=0.18$; bout number: $F(1,33) = 1.86$; $p=0.18$) nor viral expression (distance: $F(1,33)=0.29$; $p=0.59$; time spent running:

$F(1,33)=1.10$; $p = 0.31$; speed: $F(1,33) = 10.82$; $p=0.37$; bout number: $F(1,33) = 0.001$; $p=0.98$) impacted VWR architecture during the inactive cycle. All interactions were $p < 0.05$. Data points are means ± SEM.
